# Engineering functional membrane-membrane interface by InterSpy

**DOI:** 10.1101/2022.04.04.487023

**Authors:** Hossein Moghimianavval, Chintan Patel, Sonisilpa Mohapatra, Sung-Won Hwang, Tunc Kayikcioglu, Yashar Bashirzadeh, Allen P. Liu, Taekjip Ha

## Abstract

Engineering synthetic interfaces between membranes has potential applications in designing non-native cellular communication pathways and creating synthetic tissues. Here, InterSpy is introduced as a synthetic biology tool consisting of a heterodimeric protein engineered to form and maintain membrane-membrane interfaces between apposing synthetic as well as cell membranes through SpyTag/SpyCatcher interaction. Inclusion of split fluorescent protein fragments in the designed InterSpy toolkit allows tracking the formation of membrane-membrane interface and reconstitution of functional fluorescent protein in the space between apposing membranes. We first demonstrate InterSpy by testing split protein designs using a mammalian cell-free expression system. By utilizing co-translational helix insertion, cell-free synthesized InterSpy fragments are incorporated into the membrane of liposomes and supported lipid bilayers with a desired topology. Functional reconstitution of split fluorescent protein between the membranes is strictly dependent on SpyTag/SpyCatcher. Finally, since InterSpy is fully genetically encoded, the engineered system is adapted to cells and showcased. InterSpy demonstrates the power of cell-free expression systems in functional reconstitution of synthetic membrane interfaces via proximity-inducing proteins. This technology may also prove useful for synthetic biology where cell-cell contacts and communication are recreated in a controlled manner using minimal components.

## Introduction

Contacts between membranes occur naturally and ubiquitously inter- and intracellularly. Functional roles of intercellular membrane interfaces include leak-proofing endothelial layers via claudin 1 in tight junctions [1,2], creating an epithelial cell monolayer with E-cadherins [3–5], and establishing communication in the nervous system where neurons communicate with each other or innervate muscle cells [6–8]. In addition, intracellular membrane interfaces between organelles and plasma membrane known as membrane contact sites have been identified and intensely studied [9–11]. Intercellular membrane interfaces are involved in processes such as signaling [12,13], mechanosensation [14–16], and defining membrane topology [17] or tissue morphogenesis [18], whereas intracellular membrane contact sites play crucial roles in autophagy [19,20], lipid signaling and transport [21–25], and organelle trafficking [26–28].

Recent discoveries have demonstrated the distinct physicochemical properties of membrane interfaces such that while certain proteins become immobilized, others retain their 2D diffusional freedom [29]. The interface height has been shown to dictate membrane protein sorting and size-exclude large proteins [30]. While membrane contacts occur naturally, engineering a functional membrane interface is of great interest as it allows the study of protein-protein interactions that occur at such interfaces, creation of a network of cells or organelles for biomimicry of a naturally occurring phenomenon, or synthesis of an active material, to name a few.

One area where membrane interfaces is particularly important is in synaptic transmission. It has been shown that HEK293 cells can be engineered to become excitable cells and fire action potentials [31–33]. To create a network from electrically communicating excitable cells, a system that brings and maintains membranes close to each other for electrochemical exchange without allowing ion transfer [34] (e.g. through connexins) or fusion (e.g. facilitated by SNARE proteins) is required. From a therapeutic point of view, a molecular tether that preserves the distinct identity of each membrane while keeping them in close proximity can be useful. In this context, stable binding of liposomes as drug carriers to cells permits localized drug release from liposomes to selected targets with minimal toxicity to non-targeted cells. The key element in such systems is an engineered membrane interface that restricts and spatially organizes reactions or molecular release at the membrane interface.

Since protein-protein or protein-membrane interactions are required for natural inter- and intracellular interface formation, it can be envisioned that a synthetic membrane interface can also be engineered through designed protein-protein interactions. A simple strategy to artificially induce stable protein-protein interactions is by designing protein dimers that interact with high specificity and affinity. Successful dimerization based on hydrophobic interactions [35], metal affinity [36], and helix-helix interaction [37], for example, have been shown to be useful for designing protein-protein interaction probes, supramolecular assemblies, or even inducing membrane fusion. Another example of a highly specific chemically inducible dimerization pair is FKBP-FRB that has been used recently to induce formation of membrane contact sites [38,39]. The interaction between FKBP and FRB, however, is non-covalent and requires rapamycin for dimerization. A covalent interaction between two fragments creates an irreversible dimer and would be desirable for creating membrane interfaces.

Covalent bond formation between two protein fragments can be engineered to occur through cysteine residues via disulfide bridges [40], incorporation of reactive non-canonical amino acids [41,42], or spontaneous isopeptide [43,44], ester [45], or thioester [46] covalent bond formation. Of these, cystine bonds are sensitive to the redox environment and might need site-specific mutation for creating a disulfide bridge, and non-canonical amino acid incorporation requires further steps in protein expression and a careful choice for the site of non-canonical amino acid incorporation. Spontaneous isopeptide bond formation between an aspartic acid residue (Asp 117) on a peptide tag called SpyTag and a lysine residue (Lys31) on a protein called SpyCatcher derived from CnaB2 domain of FbaB protein from *Streptococcus pyogenes* was described by Zakeri *et al* [43]. This system has been used for bacterial surface display of recombinant proteins [47], protein purification [48], biopolymer and hydrogel production [49], single-molecule manipulation using optical tweezers [50], and SARS-CoV-2 vaccine development [51]. SpyTag-SpyCatcher-mediated dimerization and functional reconstitution of split proteins have found applications such as split Cre reconstitution to monitor simultaneous gene expression [52] and split fluorescent protein reconstitution for biomarker development [53]. The fact that Spy chemistry results in covalent conjugation and prior demonstrations of its potential in mediating protein reconstitution make SpyTag/SpyCatcher a promising candidate for creating, maintaining, and functionalizing membrane interfaces.

Here we introduce InterSpy, a heterodimeric protein engineered to form and maintain membrane interfaces between apposing synthetic as well as cell membranes through SpyTag/SpyCatcher interaction. Each constituent member of InterSpy consists of a split fluorescent protein component and a SpyTag or SpyCatcher domain to ascertain covalent bond formation. Additionally, InterSpy fragments are engineered to possess one-pass transmembrane domains that facilitate protein incorporation into the membrane of liposomes or supported lipid bilayers. We demonstrate that in a cell-free environment, InterSpy forms membrane interfaces and facilitates local reconstitution of the split fluorescent protein at the interface through the SpyTag/SpyCatcher interaction. We apply the InterSpy design to a cellular system and demonstrate its functionality in forming and maintaining cell membrane-membrane interfaces. Our findings highlight the potential of InterSpy as a synthetic biology tool that can aid study of protein interactions in membrane interfacial regions, creation of synthetic communication pathways, or generation of artificial tissues.

## Results

### Engineering InterSpy components for protein reconstitution in cell-free and in cellular environments

We were interested in repurposing the SpyTag and SpyCatcher peptide-protein pair [43] to facilitate split protein reconstitution in cell-free as well as in cellular environments. A split fluorescent protein allows routine fluorescence imaging to be used to visualize the reconstitution of a functionally active protein. We fused split fragments of superfolder cherry 2 (called sfCherry hereafter) [54] with SpyTag and SpyCatcher. The split sfCherry fragments are sfCherry_1-10_ containing the first 10 β -sheet strands from the full sfCherry protein and sfCherry_11_ which contains the 11^th^ β -sheet strand [54]. Feng *et al*. showed that when sfCherry_11_ is bound to histone 2B (H2B), fusing SpyTag to the N-terminus of sfCherry_11_ and SpyCatcher to the C-terminus of sfCherry_1-10_ increases the brightness of sfCherry in the nucleus [53]. We adapted the existing split protein system and asked whether SpyTag-SpyCatcher interaction can mediate reconstitution of split sfCherry components in both cell-free and cellular environments. We envisioned this engineered SpyTag-SpyCatcher toolkit (**Figure 1**) could function as a membrane-membrane interface tether and facilitate split protein reconstitution in diverse environments.

**Figure 1:**
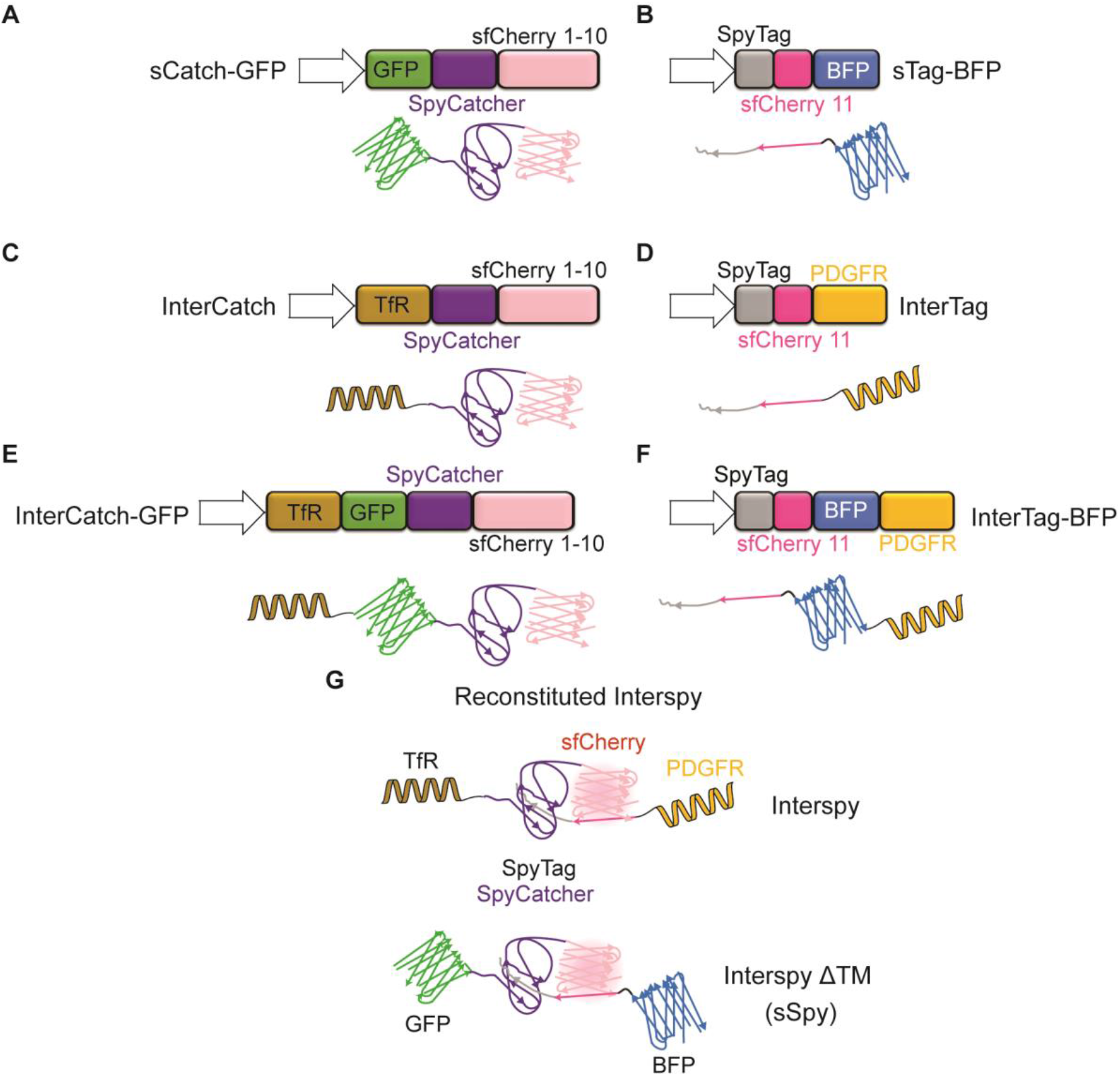
Schematic illustration of genetic constructs (top panels) and their corresponding protein variants (bottom panels) used in this work (**A-F**). **A)** sCatch-GFP: Split sfCherry_1-10_ fused to SpyCatcher and sfGFP without a transmembrane domain. **B)** sTag-BFP: Split sfCherry_11_ fused to SpyTag and TagBFP without a transmembrane domain. **C)** InterCatch: Split sfCherry_1-10_ fused to SpyCatcher and single-pass transmembrane domain of transferrin receptor (TfR). **D)** InterTag: Split sfCherry_11_ fused to SpyTag and single-pass transmembrane domain of platelet derived growth factor receptor (PDGFR). **E)** InterCatch-GFP: Split sfCherry_1-10_ fused to SpyCatcher, sfGFP and TfR transmembrane domain. **F)** InterTag-BFP: Split sfCherry_11_ fused to SpyTag, TagBFP and PDGFR transmembrane domain. **(G)** SpyTag/SpyCatcher assisted reconstitution of fluorescent sfCherry using transmembrane and soluble versions of InterSpy.

To investigate reconstitution of protein dimerization and complementation in a cell-free environment, we designed two soluble protein constructs called sCatch and sTag, where ‘s’ stands for soluble, that express SpyCatcher-sfCherry_1-10_ and SpyTag-sfCherry_11_, respectively. The corresponding fluorescent variants were generated through fusion of sfGFP (referred to as GFP, hereafter) and TagBFP (referred to as BFP, hereafter) to sCatch and sTag, respectively, resulting in sCatch-GFP (**Figure 1A**) and sTag-BFP. (**Figure 1B**). For cell surface expression or reconstitution of sCatch and sTag on supported lipid bilayers, we generated transmembrane versions of sCatch and sTag, called InterCatch (**Figure 1C**) and InterTag (**Figure 1D**), respectively, and their fluorescently tagged respective versions InterCatch-GFP (**Figure 1E**) and InterTag-BFP (**Figure 1F**). The engineered system with all necessary components that enable successful protein reconstitution in membrane interfaces is referred to as “InterSpy” (**Figure 1G**, top Panel). The soluble version of InterSpy (devoid of transmembrane domains) with fluorescent protein fusions is called “sSpy” (**Figure 1G**, bottom panel).

### Establishing InterSpy design in a cell-free expression system

We began with reconstituting sSpy since it does not have potential insolubility problems that might cause protein aggregation that interferes with dimer formation and sfCherry reconstitution. In order to reconstitute the sSpy system in a cell-free environment, we set out to investigate the functionality of soluble domains of each component. To show that *de novo* synthesized proteins form a dimer and correctly fold to reconstitute fluorescent sfCherry, we used a HeLa-based cell-free expression (CFE) system to synthesize the proteins sCatch and sTag (**Figure 2A**). We fused sCatch to GFP and sTag to BFP to monitor the kinetics of the CFE reactions (**Figure 2B**). We found that cell-free synthesized proteins form a heterodimer mediated by the isopeptide bond formation between the SpyTag and SpyCatcher domains. After completion of separate CFE reactions, by mixing two CFE reaction products, we detected dimers by using SDS-PAGE (**Figure 2C**, left panel). In contrast, removal of SpyTag and SpyCatcher domains from sTag-BFP and sCatch-GFP, respectively, prevented dimer formation (**Figure S1A**). These constructs were named sTag-BFP-ΔTag and sCatch-GFP-ΔCatcher, respectively. To confirm that sfCherry fluorescence reconstitution follows dimer formation, we monitored sfCherry signal and found that after almost 1 hour, the sfCherry signal started to rise even though the isopeptide bond formation is reported to be on minutes time scale [55] (**Figure 2C**, right panel). Deletion of SpyTag/SpyCatcher domains prevented reconstitution of sfCherry fluorescence (**Figure 2C**, right panel), further highlighting the importance of molecular proximity induced by the SpyTag/SpyCatcher conjugation system, in the absence of which the split fragments did not reconstitute to a detectable level.

**Figure 2:**
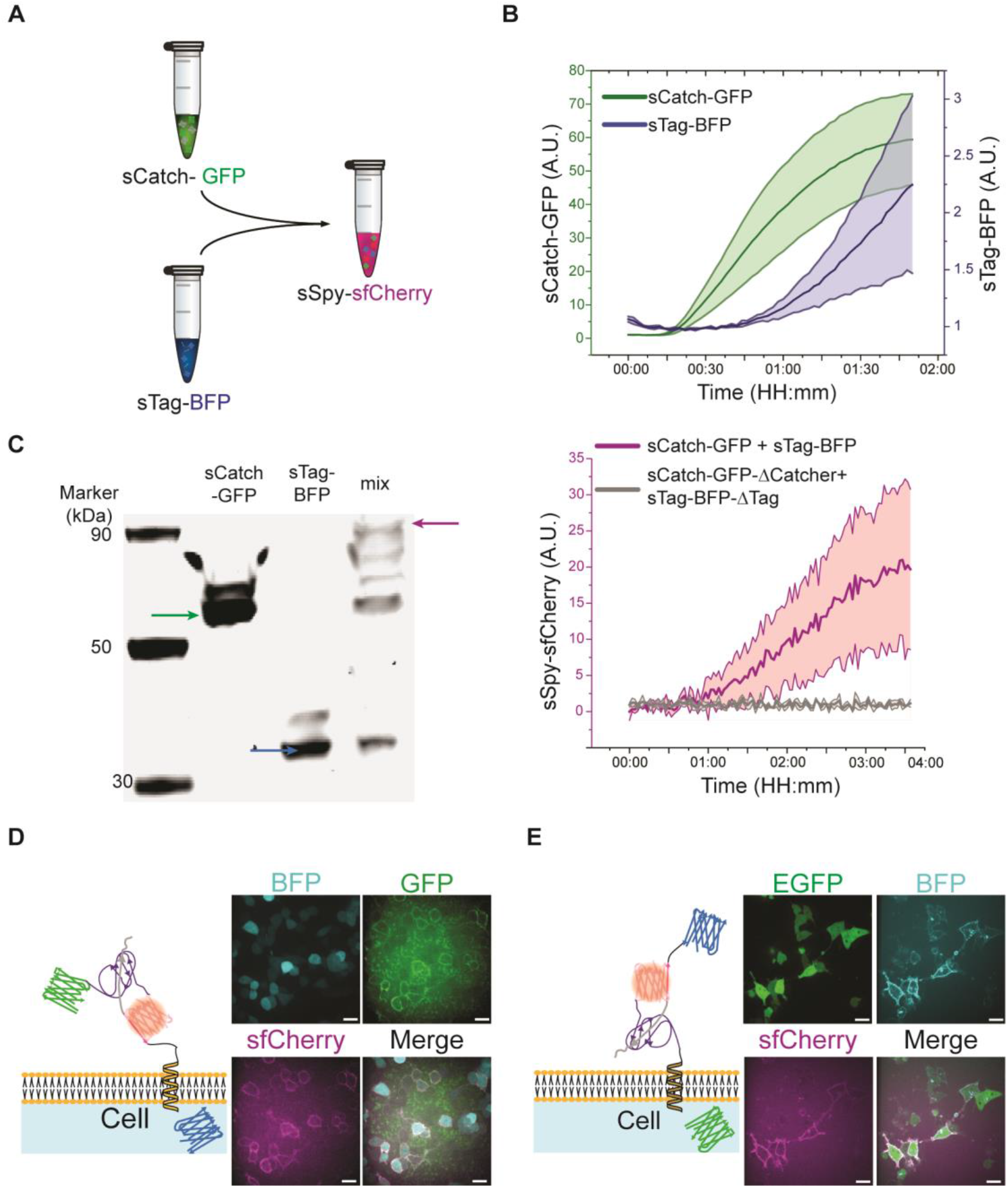
sTag-BFP and sCatch-GFP expressed through the CFE system are functionally reconstituted *in vitro*. **A)** Schematic representation of *in vitro* reconstitution of sfCherry through interaction of sCatch-GFP and sTag-BFP. sCatch-GFP and sTag-BFP are synthesized in a cell-free system via coupled transcription-translation. **B)** Translation of sCatch-GFP (green) and sTag-BFP (blue) during cell-free protein synthesis reactions were tracked through fluorescence readouts. **C)** Left: In-gel fluorescence imaging of ladder (Lane 1), sCatch-GFP (Lane 2), sTag-BFP (Lane 3), and a mixture of sTag-BFP and sCatch-GFP (Lane 4). The signals of GFP and BODIPY-FL lysine indicated protein bands for full-length expression of sCatch-GFP (Lane 2, green arrow) and sTag-BFP (Lane 3, blue arrow), respectively. When two proteins were mixed, a protein band around 90 kDa appeared (Lane 4, red arrow). Right: (Red) Fluorescence readout of reconstituted sfCherry upon mixing sTag-BFP and sCatch-GFP. (Gray) Fluorescence readout from a control experiment where sCatch-GFP-ΔCatcher and sTag-BFP-ΔTag were mixed. **D)** Representative confocal images of reconstituted sfCherry (magenta) through addition of sCatch-GFP (green) to HEK293T cells displaying InterTag on their plasma membrane identified by BFP (cyan) cytoplasmic expression. Also shown is the schematic of reconstituted sfCherry through interaction of SpyTag/SpyCatcher at the membrane. **E)** Representative confocal images of sfCherry (magenta) reconstitution by addition of cell-free synthesized sTag-BFP (cyan) to HEK293T cells expressing InterCatch at the membrane identified by EGFP (green) cytoplasmic expression. Scale bars: 20 μm.

We next asked if sfCherry reconstitution can occur via SpyTag-SpyCatcher even when one partner possesses less diffusional freedom. We fused the N-terminus of sCatch to single-pass transmembrane domain of transferrin receptor (TfR) and C-terminus of sTag to the single-pass transmembrane of platelet-derived growth factor receptor beta (PDGFRβ) to make InterCatch and InterTag (**Figures 1C and 1D**), respectively, and expressed them separately in HEK293T cells. To help identify cells that express InterTag and InterCatch, the constructs included BFP and EGFP, respectively, following a P2A self-cleaving peptide such that cells that expressed InterTag had soluble BFP and cells that expressed InterCatch had soluble EGFP. As expected, we observed sfCherry reconstitution on the surface of InterTag expressing cells when the cells were mixed with sCatch-GFP produced by CFE (**Figure 2D**). Similarly, addition of cell-free synthesized sTag-BFP to cells expressing InterCatch led to cell surface reconstitution of sfCherry (**Figure 2E**). Additionally, we found that deletion of SpyCatcher domain from sCatch-GFP eliminated sfCherry reconstitution (**Figure S1B**). Together, these results highlight the importance of SpyTag-SpyCatcher covalent bond formation prior to sfCherry reconstitution in CFE reactions *in vitro* and on cell membranes.

### Reconstitution of InterSpy on supported lipid bilayers by direct insertion of cell-free expressed membrane proteins

To expand the applicability of the reconstituted InterSpy system to a cell-free environment, we utilized the ability of the HeLa CFE system to co-translationally translocate transmembrane proteins into lipid bilayers. We used supported lipid bilayers with excess reservoir (SUPER) templates that have been used in previous synthetic biology applications to demonstrate lipid bilayer incorporation of cell-free synthesized proteins [56,57]. SUPER templates are generated via spontaneous rupture of small unilamellar vesicles (SUVs) on the silica surface of spherical beads [58]. Unless mentioned otherwise, we used 5 μm beads.

We fused the N-terminus of sCatch-GFP with TfR transmembrane domain to create InterCatch-GFP and C-terminus of sTag-BFP with PDGFRβ transmembrane domain to create InterTag-BFP (**Figures 1E** and **1F**). We relied on liposome-assisted translocation of alpha-helical transmembrane domains during translation into supplied SUVs and used the resulting protein-bound SUVs to generate SUPER templates (**Figure 3A**). Using this strategy, we showed insertion of InterTag-BFP onto the membrane of SUPER templates (**Figure 3B**). When we mixed SUPER templates harboring InterTag-BFP on their membranes with cell-free produced sCatch-GFP, we observed reconstitution of functional sfCherry localized to the membrane of SUPER templates (**Figure 3B**). This not only illustrates the ability of our system to reconstitute the split protein localized on the membrane of SUPER templates, but also allows us to ascertain the orientation of the reconstituted membrane protein. Intuitively, hydrophobic interactions require that a single transmembrane domain inserts into the lipid bilayer during translation while the rest of the protein remains soluble in the outer, exoplasmic environment. Additionally, an SUV can rupture on a glass surface such that the inner leaflet contacts the glass surface, leaving the extravesicular domain on the solution side [59]. In our case, we conclude that at least some of the transmembrane proteins have an outward-facing topology in which the soluble domains face the outer environment of the SUPER template, referred to as exoplasmic side, and can interact with the added sCatch or sTag. Expectedly, the sfCherry fluorescence signal was not observed when sCatch-GFP-ΔCatcher was mixed with InterTag-BFP harboring SUPER templates (**Figures 3C** and **S2A**). These results confirm our previous observations where sCatch-GFP failed to reconstitute functional sfCherry in the absence of SpyCatcher on cell membrane.

**Figure 3:**
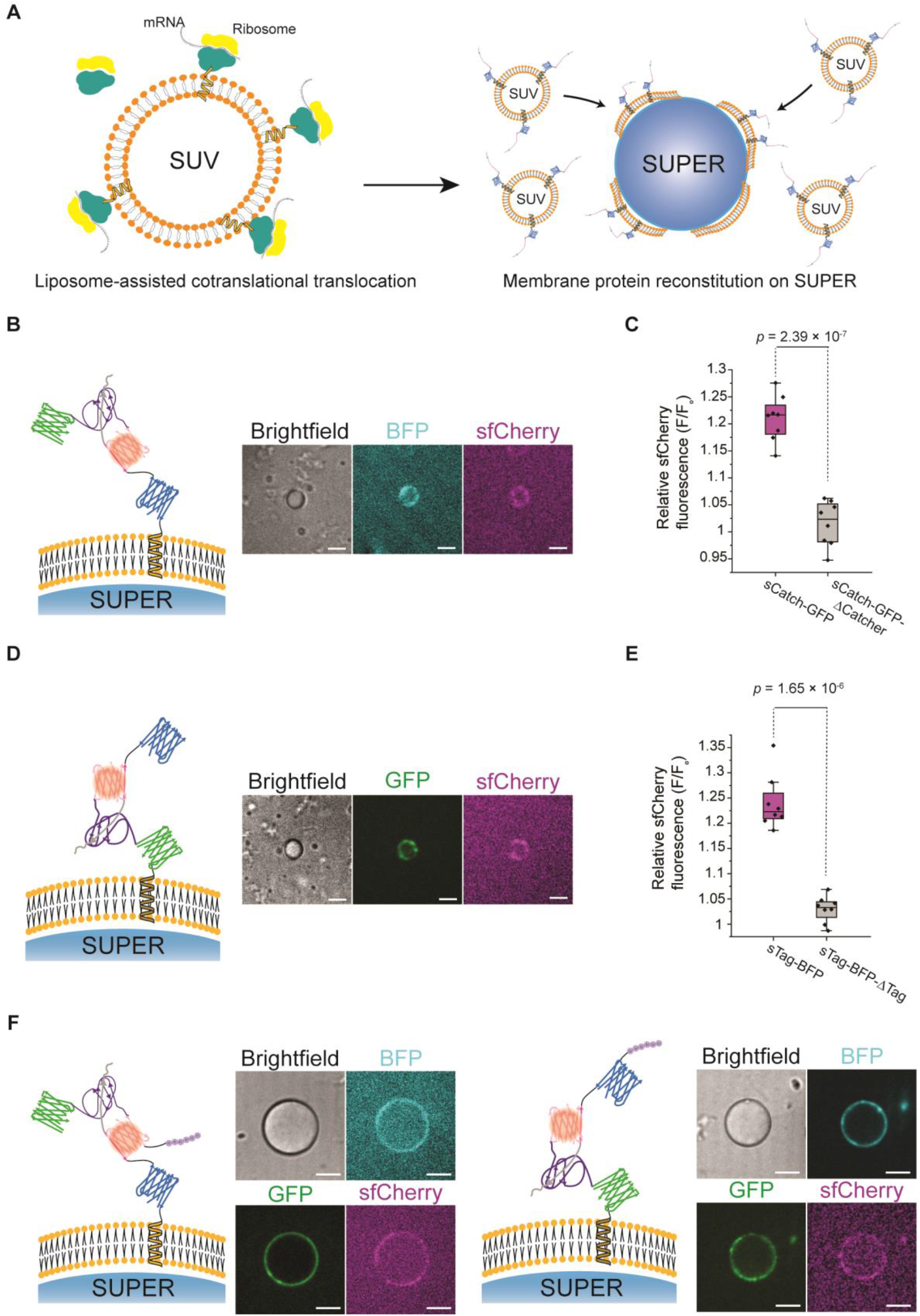
Addition of a transmembrane domain to SpyTag/SpyCatcher proteins allows mCherry reconstitution on the membrane of supported lipid bilayers. **A)** Schematic representation of the liposome-assisted translocation of membrane protein InterTag-BFP during cell-free expression (left) and membrane protein reconstitution on SUPER templates through deposition of liposomes harboring fully-translated proteins on silica beads (right). **B)** Representative confocal images of reconstitution of cell-free expressed InterTag-BFP (cyan) on the membrane of 5 μm SUPER templates and reconstitution of sfCherry (magenta) on the membrane. Scale bars: 5 μm. **C)** Box plots comparing the relative sfCherry signal on the membrane of SUPER templates in the presence (magenta) or absence (gray) of SpyCatcher domain. The data shows the average ratio of sfCherry signal on the SUPER membrane to the background signal for 30 points along the bead periphery across 8 different beads. The box represents the 25–75th percentiles, and the median is indicated. The whiskers show the minimum and maximum data points. **D)** Representative confocal images of reconstitution of sfCherry (magenta) on the membrane of 5 μm SUPER templates where InterCatch-GFP (green) was incorporated into the membrane. Scale bars: 5 μm. **E)** Box plots depicting the relative mCherry fluorescence signal on the membrane of SUPER templates in the presence (magenta) versus absence (gray) of SpyTag domain. The data shows the average ratio of mCherry signal on the SUPER membrane to the background signal for 30 points along the bead periphery in 8 different beads. **F)** Representative confocal images of bottom-up reconstitution of membrane proteins InterTag-BFP (cyan, left) and InterCatch-GFP (green, right) on the membrane of 20 μm SUPER templates and sfCherry (magenta) formation when mixed with purified sCatch-GFP (green) and sTag-BFP (cyan), respectively. Scale bars: 10 μm.

Exploiting the same strategy as described above, we showed that InterCatch-GFP incorporates into the membrane of SUPER templates with exoplasmic exposure similar to InterTag-BFP. Addition of cell-free synthesized sTag-BFP led to sfCherry reconstitution on SUPER templates (**Figure 3D**). This confirms the generality of liposome-assisted protein insertion in membrane protein reconstitution using CFE. A control experiment adding sTag-BFP-ΔTag to SUPER templates with InterCatch-GFP did not show sfCherry reconstitution (**Figures 3E** and **S2B**). Interestingly, in some cases we observed no GFP signal on the membrane of SUPER templates while we were able to detect sfCherry signal. In such cases, we detected the occurrence of fluorescence resonance energy transfer where we observed sfCherry emission by exciting GFP and a complete loss of GFP fluorescence (**Figure S3**).

We next reconstituted InterTag-BFP and InterCatch-GFP on larger, 20 μm silica beads as their sizes are similar to mammalian cells for developing cell-SUPER interface in the future. When reconstituted on 20 μm beads, sfCherry signal strength was severely diminished likely due to the low yield of low-volume CFE reactions. Consistent with this interpretation, we observed robust sfCherry fluorescence when we used soluble partners purified from bacterial expression (**Figure 3F**, see **Figures S4A** and **S4B** for controls).

### Reconstitution of protein dimerization between two apposing membranes through InterSpy

Upon verifying successful incorporation of InterCatch-GFP and InterTag-BFP with the appropriate topology for mediating functional protein reconstitution on the SUPER templates, we investigated the possibility of reconstituting the InterSpy system in the space between two apposing synthetic membranes. To create the interface between two synthetic membranes, we used SUVs and SUPER templates. We utilized liposome-assisted insertion of cell-free synthesized InterTag-BFP and InterCatch-GFP into the lipid bilayers to generate protein-coated large 20 μm SUPER templates. Additionally, we dosed the SUV membrane with a trace amount of DGS-NTA-Ni lipid and used polyhistidine and NTA-Ni affinity to bind either purified sCatch-6xHis or purified sTag-6xHis to SUVs. To avoid the possible unbinding of His-tagged proteins from NTA-Ni lipids, no competing reagent was included in the solution and the protein-bound SUVs were prepared right before the reconstitution assay.

After purifying SUVs harboring sCatch-6xHis using size exclusion chromatography, we mixed the SUVs and InterTag-BFP-containing SUPER templates and observed sfCherry reconstitution at their interface (**Figure 4A**). A control experiment omitting SpyCatcher did not show sfCherry signal, showing that even when both proteins are restricted to membranes, the sfCherry fluorescence reconstitution still requires proximity facilitated by interaction between SpyTag and SpyCatcher (**Figure 4B**). Similarly, when we reconstituted InterCatch-GFP on the SUPER templates and added sTag-BFP-6xHis bound to SUVs, we detected sfCherry formation at the membrane interface mediated by SpyTag/SpyCatcher (**Figures 4C and D**).

**Figure 4:**
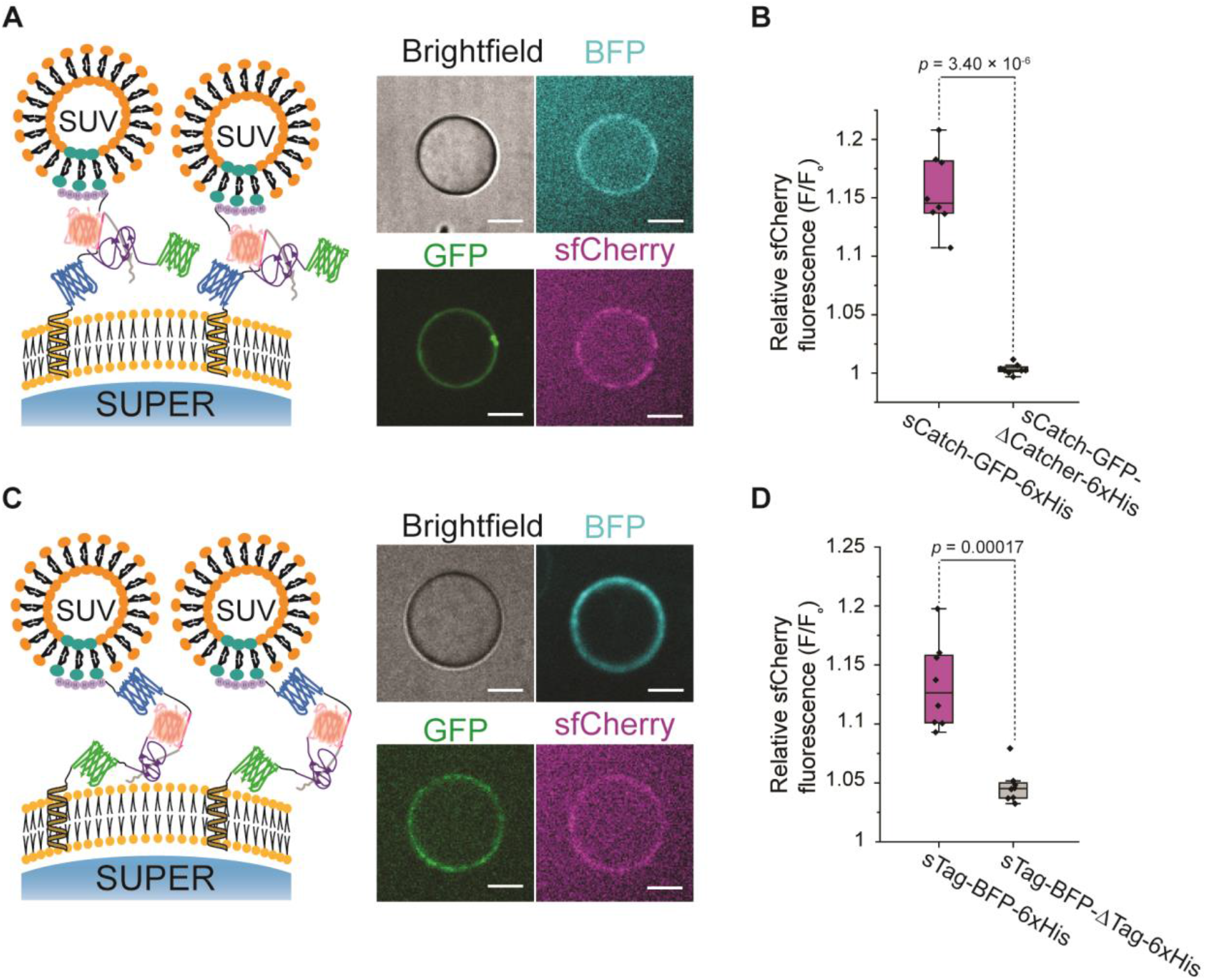
Cell-free reconstituted fluorescent sfCherry between two apposing membranes. **A)** Representative confocal images of reconstituted sfCherry (magenta) that fluoresces in the space between SUVs harboring sCatch-GFP (green) and SUPER templates displaying InterTag-BFP (cyan). Scale bars: 10 μm. **B)** Box plots comparing the relative sfCherry signal in the presence (magenta) or absence (gray) of SpyCatcher domain. The ratios of sfCherry signal on the SUPER membrane to the background signal were averaged over 30 points along the bead periphery to account for non-uniform distribution of fluorescence over the bead surface. F/F_0_ is the mean value of the ratios, averaged across 8 different beads. **C)** Representative confocal images of reconstituted sfCherry (magenta) that fluoresces in the space between SUVs harboring sTag-BFP (cyan) and SUPER templates displaying InterCatch-GFP (green). Scale bars: 10 μm. **D)** Box plots comparing the relative sfCherry signal in the presence (magenta) or absence (gray) of SpyTag domain. The ratios of sfCherry signal on the SUPER membrane to the background signal were averaged over 30 points along the bead periphery to account for non-uniform distribution of fluorescence over the bead surface. F/F_0_ is the mean value of the ratios, averaged across 8 different beads.

We cannot rule out the possibility of membrane fusion between SUVs and SUPERs that caused sfCherry formation via cis-interaction. However, given the composition of SUPER templates and SUVs that lack any fusogenic lipids and the calcium- and magnesium-free exoplasmic solution (1x PBS), these interactions are not expected to occur [60].

Consistent with our observation in bulk experiments where sfCherry fluorescence appeared long after SpyTag-SpyCatcher bond formation (**Figure 2C**, right panel), we noticed that SUVs harboring sTag-BFP-6xHis localized to InterCatch-GFP on SUPER templates only a few minutes after addition of SUVs due to the SpyTag-SpyCatcher interaction, but sfCherry formation and fluorescence occurred after a few hours (**Figure S5**). We note that reconstitution of sfCherry in a membrane-membrane interface with the constituent membrane proteins incorporated into their corresponding lipid bilayers with alpha helical transmembrane domain was not feasible due to the low yield of protein-bound SUVs after CFE reaction and size exclusion chromatography.

Additionally, to demonstrate that InterSpy functions under a variety of conditions and can be used for diverse applications, we used purified sCatch-GFP-6xHis and sTag-BFP-6xHis to reconstitute the InterSpy between DGS-NTA-Ni-containing SUPER templates and SUVs. Under this condition, sfCherry reconstitution was observed with high efficiency (**Figure S6A**). Even though using purified components allowed us to use higher concentrations of sCatch-GFP and sTag-BFP, the sfCherry reconstitution remained dependent on the presence of both SpyTag and SpyCatcher domains (**Figure S6B**). These results indicate that in the absence of SpyTag/SpyCatcher interaction, increasing the concentration of split protein fragments does not necessarily lead to reconstitution and corroborated the essential role of isopeptide bond formation for inducing proximity and local immobilization of split protein fragments. Along with our earlier findings showing the slow formation of sfCherry (**Figure 2C** right panel and **Figure S5**), we conclude that the sfCherry fragments may have such a low affinity that self-assembly of its split fragments does not occur to a detectable degree without SpyTag/SpyCatcher’s induced proximity.

### InterSpy for intercellular protein complementation

We showed earlier that InterTag or InterCatch construct expressed on the surface of cell membranes facilitated sfCherry formation via isopeptide bond formation when the complementary fragment was synthesized by a CFE system in a soluble form (**Figures 2D** and **2E**). We next wanted to reconstitute the InterSpy in membrane-membrane interfaces in a fully cellular environment.

In our preliminary experiments, we co-cultured HEK293T cell lines that were transiently expressing either InterTag-BFP or InterCatch-GFP. We observed that successful expression of InterTag-BFP and InterCatch-GFP allowed for reconstitution of the split sfCherry components at cell-cell contact regions with 97% yield (**Figure S7A**). We then investigated whether the presence of the fused fluorescent proteins (i.e., GFP and BFP) as part of the InterTag-BFP and InterCatch-GFP, respectively, is dispensable for sfCherry reconstitution at intercellular junctions. We generated two stable HEK293T cell lines that express InterTag and InterCatch constructs individually (**Figures 1C** and **1D**), with cytosolic expression of BFP and EGFP, respectively.

We imaged the co-cultured cell line pair stably expressing InterTag (identified through BFP fluorescence) and InterCatch (identified through EGFP fluorescence) and observed reconstituted sfCherry fluorescence signal (**Figure 5A**, left panel). As expected, the sfCherry fluorescence was restricted only to intercellular junctions between two cells, one expressing InterTag and the other expressing InterCatch. Even though InterTag-InterCatch cell pairs formed membrane-membrane contact, the vast majority (∼ 95%) of these intermembrane junctions did not show sfCherry reconstitution (**Figure 5A**, right panel). The significant difference in the sfCherry reconstitution efficiency at cellular junctions of cell pairs expressing InterTag-BFP/InterCatch-GFP vs. those expressing InterTag/InterCatch suggested that the removal of the respective fused fluorescent protein led to the observed dramatic decrease in the reconstitution efficiency from ∼97 % to ∼5 % (**Figure S7B**). We reasoned that the SpyTag and SpyCatcher domains alone may not be able to span the distance between the cell membranes to allow interaction between the split constituents of sfCherry.

**Figure 5:**
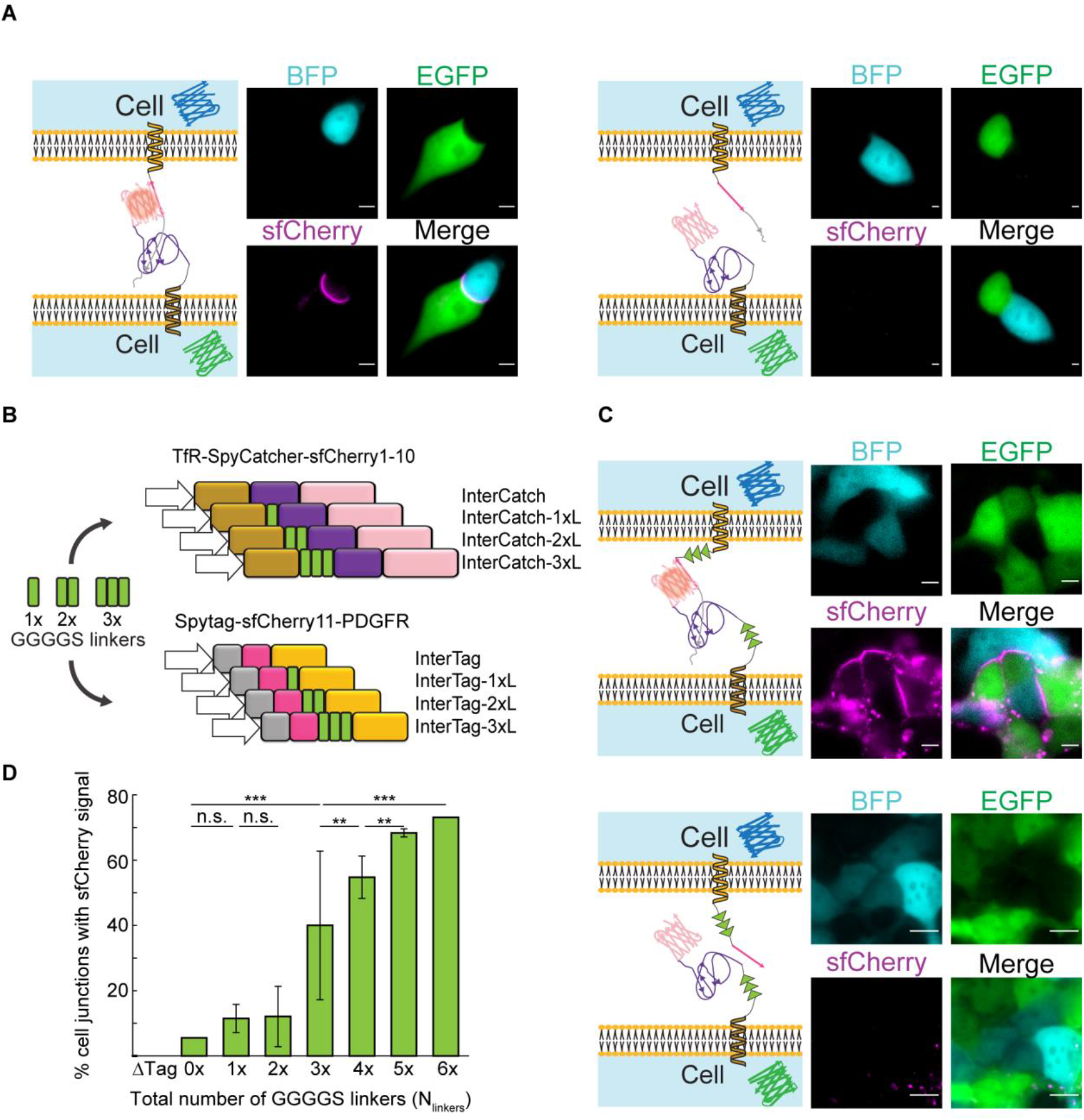
Effect of number of GGGGS linkers in the InterSpy system on the efficiency of reconstitution of sfCherry fluorescence. **A)** Left: Representative single cell image of InterSpy-0xL (sfCherry, magenta) reconstituted through InterTag-0xL (BFP, cyan) and InterCatch-0xL (EGFP, green). Right: A representative single cell image of unsuccessful reconstitution of InterSpy-0xL (sfCherry). Corresponding representative images of cells expressing InterTag-0xL (BFP, cyan) and InterCatch-0xL (EGFP, green) are also shown. **B)** Schematic representation of a mini-library of InterTag and InterCatch constituting different numbers of GGGGS linkers. **C)** Top: Representative cell cohort image of reconstituted InterSpy-6xL (sfCherry, magenta) with InterTag-3xL (BFP, cyan) and InterCatch-3xL (EGFP, green). Bottom: Representative cell cohort image showing lack of sfCherry signal upon interaction of InterTag-3xL-ΔTag (BFP, cyan) and InterCatch-3xL (EGFP, green). **D)** Plot showing percent of cell-cell junctions with reconstituted sfCherry signal vs. total number of intercellular GGGGS linkers (N_linkers_). The error bars represent the standard deviation across experiments performed for co-cultured cell line pairs expressing the same total number of GGGGS linkers. E.g. co-cultured cell pairs expressing InterTag-0xL + InterCatch-3xL, InterTag-1xL + InterCatch-2xL, InterTag-2xL + InterCatch-1xL, and InterTag-3xL + InterCatch-0xL

To compensate for the absence of fused fluorescent proteins in the InterTag and InterCatch constructs and to assist with the establishment of the interaction between SpyTag and SpyCatcher, we introduced flexible glycine-serine (GGGGS) linkers into the InterTag and InterCatch constructs. We generated a mini library consisting of a total of 8 plasmid constructs, introducing zero, one, two, or three repeats of GGGGS linkers between the transmembrane protein and the SpyTag/SpyCatcher component of the protein (**Figure 5B**). We established stable cell lines that express InterTag-#xL and InterCatch-#xL where # denotes the number of GGGGS linkers integrated into the InterTag/InterCatch construct and ranges from 0 to 3 and L denotes linker. We then tested all 16 permutations of co-culture pairings between the InterTag-#xL and InterCatch-#xL expressing stable cell lines such that the total number of GGGGS linkers, *N*_linkers_, that span the intercellular region varied from 0 to 6. For *N*_linkers_ = 6, we saw a dramatic increase in reconstitution efficiency to ∼70 % (**Figure 5C**, top panel, **Figure 5D**) suggesting that intercellular reconstitution is affected by the steric effect between InterTag and InterCatch which can be regulated by tuning the linker lengths. We further confirmed the need for the Spy system using the InterTag-ΔTag constructs that showed 0% reconstitution efficiency (**Figure 5C**, bottom panel). Interestingly, we noticed sfCherry fluorescent puncta inside some GGGGS-linker containing-InterSpy cells, the reasons for which were not entirely clear to us.

The reconstitution efficiency increased with increasing *N*_linkers_. When *N*_linkers_ increased from 2 to 3, there was noticeable jump from ∼10% to ∼40% (**Figure 5D**) (see **Table S1** for the percentage of each combination). This suggested that the distance between cell-cell membrane is covered by inclusion of at least 3 GGGGS linkers in our engineered InterSpy system. Additional linkers still helped because with *N*_linkers_ = 6, the efficiency reached ∼70%. Altogether, these results demonstrate the minimum distance between apposing membranes must be accounted for in designing intercellular split protein reconstitution.

## Discussion

We reconstituted membrane-membrane interfaces with split protein activity mediated by SpyTag/SpyCatcher interaction in both cell-free and cellular environments using the InterSpy toolkit. We demonstrated cell-free synthesis of split protein fragments in soluble or transmembrane forms with appropriate folding that led to split protein reconstitution on cell membranes, supported lipid bilayers, and in membrane interfaces. Finally, we adapted InterSpy to a cellular system and used it to reconstitute a split fluorescent protein in intercellular interfaces.

Our results illustrated the previously unexplored power of CFE systems in aiding protein engineers for rapidly assessing the functionality of their designed proteins. We demonstrated the importance of SpyTag/SpyCatcher chemistry in inducing local proximity for split protein reconstitution. This knowledge, however, was readily provided from the starting point when cell-free synthesized soluble proteins lacking SpyTag/SpyCatcher domains did not show sfCherry signal. This example indicates how CFE systems can be utilized to rapidly evaluate the success of protein engineering by circumventing time-consuming and difficult protein expression and purification steps in a routine protein design workflow. In addition, we introduced an approach to reconstitute cell-free synthesized membrane proteins onto large protein-harboring SUPER templates. Even though we used this method to reconstitute single-pass membrane proteins, the co-translational translocation can theoretically occur while synthesizing more complex multi-pass, difficult-to-express, membrane proteins, the practical demonstration of which awaits further studies. As we demonstrated in this work, the reconstituted protein topology can be tested by inserting SpyTag domain in a presumable exoplasmic terminus or loop. Thus, introducing a fluorophore-conjugated SpyCatcher protein to SpyTag domain inserted at various locations in a membrane protein can be used to test the reconstituted protein topology. When reconstituting InterTag-BFP and InterCatch-GFP, we noted occasional occurrences of puncta formation on the periphery of SUPER templates that suggested possible microdomain formation due to the hydrophobic interactions between transmembrane domains buried inside the lipid bilayers (**Figures 3** and **4**). Such phenomena have been previously reported in similar CFE-mediated membrane protein reconstitution studies [56,57] and requires additional work in the future to investigate possible lipid-protein or protein-protein interactions.

By adapting InterSpy to HEK293T cells, we confirmed the activation of the split protein in intercellular membrane interfaces. We showed the steric effects on split protein reconstitution by varying the size of flexible linkers that led to an increase in split protein reconstitution efficiency. These results highlighted the importance of considering the physical properties of the gap between two membranes in a cell junction. In principle, it is possible to adapt these constructs in a wide variety of cell lines. Certainly, the membrane-membrane distance, which may depend on the cell type, can be tuned by changing the linker size or its rigidity and can be used to drive size-dependent protein segregation or control the membrane distance for resonance energy transfer applications mediated via either fluorescence or bioluminescence.

Our findings open up the possibility of utilizing InterSpy for applications beyond reconstitution of a fluorescent protein. There are several existing engineered protein systems such as NanoBiT [61] for bioluminescence applications (e.g. BRET), iLID [62] for light induced dimerization, and split TEV [63] for proximity-dependent chemical activation and signaling. Together with orthogonal bio-conjugation systems that are derived from other bacterial species or SpyTag/Catcher pair such as SnoopTag/Catcher [64], SdyTag/Catcher [65], and BLISS which is a light-induced SpyTag/Catcher system [66], one can imagine numerous possibilities of artificial interface designs for various applications in innovating novel cellular communication pathways.

Although we did not explore multi-cellular systems, it will be of great interest to engineer cellular networks that communicate with each other through InterSpy reconstitution system. For example, one can imagine a “synthetic synapse” in which electrical signal is transduced from the presynaptic cell by localized reconstitution and activation of NanoBiT in cellular junctions that activates light-gated ion channels such as channelrhodopsins on the postsynaptic cell, leading towards creation of an excitable tissue. Also, since we demonstrated InterSpy functionality in a cell-free system, it can be utilized in minimal systems to recreate adhesion and communication between synthetic cells [67] or drive spatial organization of proteins [68,69]. Alternatively, synthetic cells can be adhered to natural cells for therapeutic purposes such as drug delivery or to mimic processes such as phagocytosis.

## Conclusion

We introduced InterSpy, a SpyTag-SpyCatcher dimerization system for split protein reconstitution in membrane interfaces. Different versions of InterSpy constructs induced membrane interface formation and split protein reconstitution in both cell-free and in cellular environments. Cell-free reconstitution of InterSpy unveils opportunities to decorate the membrane of synthetic cells with active molecules, thus giving them more potential to be used as drug carriers or artificial immune cells. As a synthetic biology tool, InterSpy promises reconstitution of more sophisticated processes in cellular, cell-free, and hybrid interfacial regions that can be used for bottom-up study of biological processes, biomimicry of cell adhesion and communication, and creating synthetic tissues or active materials.

## Experimental Section

### Cloning and preparing DNA constructs

DNA plasmids with gene sequences encoding for SpyTag003, sfCherry_11_, SpyCatcher003, and sfCherry_1-10_were purchased from Addgene (**Table S2**). The target gene sequences were PCR amplified from the purchased plasmids using either Phusion (NEB, M0531S) or Q5 High Fidelity (NEB, M0492S) master mixes. The primers used are specified in **Table S3**. The gene fragments were assembled using Gibson or Golden Gate assembly to generate the plasmids used in this study, as described in **Table S3**.

We generated a plasmid expressing the translationally fused protein TfR-sfGFP-SpyCatcher003-sfCherry_1-10_ (InterCatch-GFP) by inserting sfCherry_1-10_ gene sequence from pcDNA3.1(+)_SpyCatcher-6aa-sfCherry_1-10_ (Feng *et al*. [53], Addgene #117484) into the TfR-sfGFP-myc tag-SpyCatcher003 plasmid (Keeble *et al*. [55]. Addgene #133451) through Gibson Assembly (primers listed in **Table S3**). We then removed the sfGFP from the fusion to generate InterCatch. Similarly, we generated the complement InterTag-BFP by inserting the sequence encoding for SpyTag002-sfCherry_11_ –TagBFP amplified from pSFFV-SpyTag-sfCherry2(11)-TagBFP (Feng *et al*. [53] Addgene #117485) into the membrane expression vector pDisplay (Addgene #34842) and replacing the SpyTag002 sequence with SpyTag003 (Keeble *et al*. [55] Addgene #133452) using Gibson Assembly. Then, we removed TagBFP from InterTag-BFP to generate the InterTag construct.

For cell-free protein synthesis, we used the pT7CFE1-6xHis-HA vector from Thermo Fisher Scientific as the backbone. We cloned TfR-sfGFP-SpyCatcher003-sfCherry_1-10_and sfGFP-SpyCatcher003-sfCherry_1-10_ into the above vector to code for InterCatch-GFP and sCatch-GFP, respectively. Similarly, we generated InterTag-BFP and sTag-BFP by cloning PDGFRβ-TagBFP-sfCherry_11_-SpyTag003 and TagBFP-sfCherry_11_-SpyTag003 into the pT7CFE1 vector, respectively. In order to retain the hydrophobicity of the C-terminal region of InterTag and the N-terminal region of InterCatch, we removed 6xHis and HA tags from pT7CFE1 vector.

For bacterial expression and purification of sCatch-6xHis and sTag-6xHis, sequences encoding for sfGFP-SpyCatcher-sfCherry_1-10_ and SpyTag-sfCherry_11_-TagBFP were cloned into a pET28b vector (generous gift from Tobias Pirzer, Technical University of Munich, Germany) such that both proteins had a 6xHis tag in their C-termini.

For stable cell expression in HEK293T, the InterCatch and InterTag sequences were cloned into pSBbi-GP and pSBbi-BP (Kowarz *et al*. [70]. Addgene #60511 and Addgene #60512) Sleeping Beauty cassettes respectively. Synthesized DNA oligos encoding for repeating glycine-serine (GGGGS) linkers were ordered as synthesized DNA oligos (Integrated DNA technologies) in a singlet, doublet, and triplet format (where the GGGGS sequence is repeated, **Table S3**) to generate the InterCatch-#xL and InterTag-#xL variants (where # represents the number of GGGGS linkers). The oligos included SapI (GCTCTTC) restriction cut sites for cloning with Golden Gate Assembly (oligos listed in **Table S3**). The linkers were incorporated after the transmembrane domain on the extracellular side as shown in **Figure 5B**. InterTag-ΔTag was generated by cloning the InterTag-#3xL without the SpyTag003 sequence.

All cloning sequences were verified by Sanger sequencing services (Eurofins or Genewiz). The assembled DNA constructs were purified from NEB XL10 Gold or DH5α competent cells (NEBC2987H) using miniprep kits (Qiagen or E.Z.N.A.® Endo-free Plasmid DNA Mini Kit I).

### Bacterial expression and purification of sTag and sCatch

Protein expression and purification were performed by following conventional His-purification methods reported elsewhere [48]. pET28b-sTag-6xHis and pET28b-sCatch-6xHis constructs were transformed into BL21-RIPL cells. Single colonies were picked from agarose plates and grown in 5 mL LB broth supplemented with 50 μg/mL kanamycin overnight at 37 °C shaking at 220 rpm in an orbital shaker. Next, the culture was diluted into 1 L of LB broth supplemented with 0.8% w/v glucose and 50 μg/mL kanamycin and was grown at 37 °C while shaking at 220 rpm until the A_600_ reached 0.5-0.6. The culture was then induced with 0.42 mM isopropyl β-D-1-thiogalactopyranoside (IPTG) and incubated at 30 °C with constant shaking at 200 rpm for 4-5 hours. The culture was then pelleted through centrifugation at 5000 g for 10 min. The pellet was resuspended in 30 mL of lysis buffer containing 50 mM Tris-HCl (pH 7.4), 300 mM NaCl, and 1 mM phenylmethylsulfonyl fluoride (PMSF). The resuspended bacteria were lysed using a sonicator (Branson Sonifier 450) and the lysate was centrifuged at 30,000 g for 25 min. The supernatant was then run through an equilibrated His-trap (GE healthcare) in an AKTA start fast protein liquid chromatography system. The column was washed with 15 column volumes of washing buffer containing 50 mM Tris-HCl (pH 7.4), 300 mM NaCl, and 50 mM imidazole. The bound protein was eluted with elution buffer composed of 50 mM Tris-HCl (pH 7.4), 300 mM NaCl, and 400 mM imidazole. The purification quality was confirmed and assessed with SDS-PAGE and the fractions with high concentrations of proteins were pooled and dialyzed against 1 L of PBS at 4 °C overnight. The protein aliquot concentration was adjusted to 0.5 mg/mL (extinction coefficients predicted by ExPasy) and stored at -80 °C until use.

### SUV preparation and size-exclusion chromatography

Vesicles of 90% 1,2-dioleoyl-sn-glycero-3-phosphocholine (DOPC) and 10% 1,2-dioleoyl-sn-glycero-3-[(N-(5-amino-1-carboxypentyl) iminodiacetic acid) succinyl] (nickel salt) (DGS-NTA-Ni) were made by following the protocol described elsewhere [71,72]. All lipids were purchased from Avanti Polar Lipids. Appropriate aliquots of lipids for a final 5 mM concentration were transferred to a clear glass vial with screw cap. Lipids were dried under a gentle stream of argon and were further incubated in a desiccator at room temperature for at least 1 h to ensure organic solvent evaporation. Then, 500 μL ultra-pure water was added to the lipid film. The mixture was vortexed until the lipid was fully dissolved, and the solution was opaque. Next, the lipid solution was passed through a 100 nm filter 11 times using an Avanti mini-extruder. The final lipid solution was stored at 4 °C and was stable for almost 2 weeks. To generate SUV-bound proteins, 50 μL solution of SUVs was mixed with 10 μL stock concentration of either sTag-6xHis or sCatch-6xHis and incubated for 5 min at room temperature. The SUV solution was then loaded on a Sepharose 4B (Sigma Aldrich) column and 200 μL fractions were eluted using PBS. A 10 μL sample of each fraction was then loaded into a 384 well conical well plate and the corresponding fluorescence intensity from either GFP or BFP was measured at 488/528 nm or 400/457 nm wavelength, respectively. The fraction containing membrane-bound protein was kept for further experiments.

### Cell-free expression reaction and direct reconstitution assembly

Thermo Fisher 1-step human coupled *in vitro* transcription-translation kit (cat# 88881) was used to synthesize proteins expressed under the T7 promoter. To express the proteins, the protocol provided by the manufacturer was followed. In short, 5 μL of HeLa lysate was mixed with 2 μL reaction mixture and 1 μL accessory proteins. The reaction volume was then brought to 10 μL by adding 10 nM DNA plasmid and ultra-pure water. In cases where direct reconstitution of InterTag or InterCatch was desired, water was replaced by 5 mM 100% DOPC SUV solution so that the final concentration of SUV in the reaction was around 1 mM. The reactions were next transferred to a 384 conical well plate and incubated in the Synergy H1 plate reader (BioTek) at 30 °C for 3-4 h. The GFP, BFP, and sfCherry signals were monitored using plate readers at 400/450 nm, 488/528 nm, and 561/625 nm excitation/emission wavelengths, respectively. For experiments that required *in vitro* translation labeling system, a 1:50 dilution of FluoroTect™ GreenLys (Promega) was added to the reaction before incubation.

### In-gel fluorescence imaging

After completion, reactions were mixed with 4x Laemmli buffer containing 10% 2-mercaptoethanol, and SDS-PAGE gels were run in a 4-20 % Bis-Tris polyacrylamide precast gel (Sigma Aldrich). Samples were not heated to avoid denaturing GFP. The GFP and GreenLys were imaged in a Sapphire Biomolecular Imager (Azure biosystems) with 488/518 nm excitation/emission wavelengths.

### Preparation of SUPER templates

After completion of CFE reactions, SUPER templates were assembled as described by Neumann *et al*. [73] with slight modifications. Briefly, 10 μL of completed CFE reaction was mixed with 10 μL 5 M NaCl, and 3.5 μL of bead solution (5 μm and 20 μm beads were purchased from Bangs laboratory and Corpuscular, respectively), corresponding to roughly 25 × 10^5^ beads for 5 μm beads and 25 × 10^3^ beads for 20 μm beads. The final volume was then brought up to 50 μL by ultra-pure water. The mixture was incubated for 30 min at room temperature with occasional gentle flicking. After incubation, 1 mL PBS was added to the solution and centrifuged for 5 min at 200 g. Next, 950 μL of supernatant was removed and the washing step was repeated twice. After the last wash, beads were resuspended in 100 μL of supernatant and were used for imaging and further assays.

### Cell culture

HEK293T (AATC, CRL-3216) cells were maintained in Dulbecco’s modified Eagle’s medium with high glucose/pyruvate (DMEM, Gibco, 11995040), supplemented with 10% (vol/vol) FBS (Gibco, 16000044), 100 μg/ml Penicillin-Streptomycin, and 0.292 mg/ml glutamine (Gibco, 10378016). All cells were grown at 37 °C and under 5% CO_2_ in a humidified incubator.

### Generation of stable cell lines

Cells were seeded in a 12 well (Falcon, 353225) chamber and grown to 60% confluency before transfection. After 24 hours, transfection was performed following the standard protocol. 100 ng of transposase-expressing helper plasmid (Mates et al. [74] Addgene #34879) was co-transfected with 1000 ng of InterTag or InterCatch, using Lipofectamine 3000 transfection reagent (Invitrogen). Two days post-transfection, the cells were replated in a 6 well (Falcon, 353224) and selected in 2 μg/ml Puromycin (STEMCELL Technologies). Every two days, the cells were replenished with 1 mL fresh media supplemented with 2 μg/ml Puromycin. After 2 weeks of selection, the resistant colonies were grown to confluency, expanded to larger flasks, counted, and frozen following standard protocols.

### Co-culturing InterSpy and InterTag expressing stable cell lines

HEK293T stable cell lines expressing variants of InterTag and InterCatch with different linkers were cultured separately in 6 well chambers (Falcon, 353046). Upon standard passaging, cells were dissociated with 200 μL of Trypsin-EDTA (0.05%, Gibco, 25300054) and seeded at a density of 1.00×10^5^ cells/mL into Nunc™ Lab-Tek™ Chambered Coverglass (155361). The following day, InterTag and InterCatch monolayers were trypsinized and resuspended into 500 μL DMEM. InterTag and InterCatch cell suspensions were mixed and transferred to an orbital shaker in the humidified incubator for 45 minutes at 100 rpm. Cell suspensions were then centrifuged and pelleted (∼200 g for 5 min). The media was aspirated and refreshed with another 500 μL of DMEM. Cells were gently resuspended via pipetting and seeded onto fibronectin-coated wells (0.5% in dPBS, Sigma-Aldrich, F1141) and incubated for 1 hour before imaging.

### Confocal imaging and image analysis

A 1:2 and 1:10 dilution of 20 μm and 5 μm beads in PBS were made, respectively, with a final volume of 50 μL, and the bead solution was transferred to a clear glass bottom 96 well plate. When observing the interaction of cell-free synthesized proteins with membrane proteins, the whole 10 μL reaction was added to the SUPER templates. In the case where purified proteins were mixed with the SUPER templates, 1 μL protein was added to the bead solution in the well. When SUV-bound proteins were supplied to the bead solution, appropriate volume was added to the SUPER templates so that the concentration of protein is similar to when purified proteins were mixed with SUPER templates. The mixture was incubated for 2-3 h in room temperature before imaging to allow reconstitution of sfCherry. Images were taken using an oil immersion UplanFL N 40 x/1.30 NA (Olympus) objective for 20 μm beads and a Plan-Apochromat 60 x/1.4 NA (Olympus) objective for 5 μm beads on an inverted microscope (Olympus IX-81) equipped with an iXON3 EMCCD camera (Andor Technology), National Instrument DAQ-MX controlled laser (Solamere Technology), and a Yokogawa CSU-X1 spinning disk confocal. Images were acquired using MetaMorph (Molecular Devices). Single images of GFP, BFP, and sfCherry proteins were taken at excitation wavelengths of 488 nm, 405 nm, and 561 nm, respectively. The intensity measurements in **Figures 3** and **4** were calculated by using ImageJ “oval profile” plug-in where an oval was drawn along the membrane of SUPER templates and intensities at 30 points, uniformly distributed along the SUPER template periphery, were measured. These intensities were then divided by the background intensity in the vicinity of the bead using the same strategy to calculate the relative sfCherry fluorescence.

### Live-cell fluorescence microscopy

Imaging was performed using Nikon Eclipse Ti-E widefield fluorescence microscope with an environmental chamber maintaining the temperature at 37 °C and an atmosphere of 5% CO2 with a 60x oil-immersion objective, unless otherwise mentioned. Images of cells expressing GFP or EGFP, BFP, and sfCherry proteins were taken at excitation wavelengths of 440 nm, 395 nm, and 550 nm, respectively.

### Statistical analysis

Image analysis data from images in **Figures 3** and **4** were exported to Microsoft Excel and two-tailed t-test with unequal variances was performed using Excel’s built-in data analysis tool to calculate p-values in **Figures 3C, 3E, 4B**, and **4D**. For the bar graph in **Figure 5D**, an R code was written and used to evaluate the association between the frequency of observation of sfCherry signal and the linker size. The code performs pairwise Fisher’s exact test to calculate p-values between each two groups, and the p-values were adjusted using Bonferroni correction with a factor of 21 since there were a total of 21 comparisons. All p-values for this graph are listed in **Table S4**. The statistical comparison between cell lines transiently transfected with either InterTag-BFP or InterCatch-GFP and cell lines expressing InterTag-0xL or InterCatch-0xL in **Figure S7B** was performed using Fisher’s exact test as well.

## Acknowledgements

The authors thank members of the building a synthetic neuron team, in particular Dr. Francois St-Pierre and Dr. Chongli Yuan, as well as members of the Ha and Liu laboratory for discussion of the manuscript. The authors acknowledge the NSF (EF1935265 to Liu and EF1934864 to Ha) and NIH (EB030031 to Liu and GM122569 to Ha) for research funding. The authors thank Dr. Tobias Pirzer of the Technical University of Munich for pET28b plasmid, Dr. Chongli Yuan of Purdue University for providing pCMV(CAT)T7-SB100 plasmid, and Dr. Cristina Mitrea for helpful advice on statistical analysis. T.H. is an investigator with the Howard Hughes Medical Institute. C.P. was funded by the Doctoral Diversity Program of Johns Hopkins Medicine. This article is subject to HHMI’s Open Access to Publications policy. HHMI lab heads have previously granted a nonexclusive CC BY 4.0 license to the public and a sublicensable license to HHMI in their research articles. Pursuant to those licenses, the author-accepted manuscript of this article can be made freely available under a CC BY 4.0 license immediately upon publication.

## Data Availability Statement

The data that support the findings of this paper are available from corresponding authors upon reasonable request.

