## Supporting Information for "Engineering functional membrane-membrane interface by InterSpy"

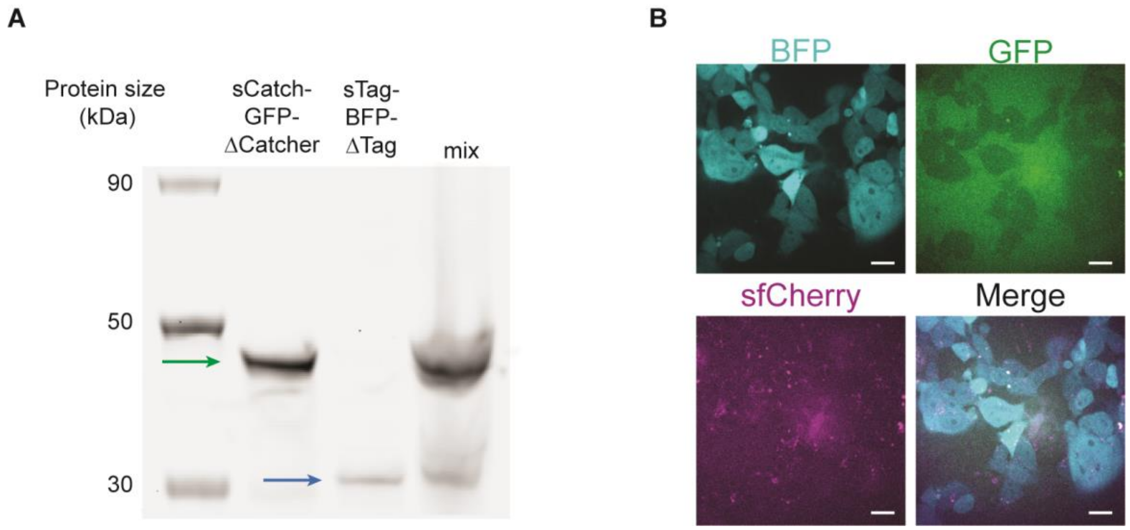


**Figure S1**: The crucial role of SpyTag/SpyCatcher domain in sfCherry reconstitution both in bulk CFE reactions and on cell membranes. **A)** In-gel fluorescence imaging of ladder (Lane 1), sCatch-GFP-ΔCatcher (Lane 2), sTag-BFP-ΔTag (Lane 3), and a mixture of sTag-BFP-ΔTag and sCatch-GFP-ΔCatcher (Lane 4). **B)** Representative confocal images of HEK293T cells expressing InterTag on their surface (identified with cytosolic expression of BFP (blue)) mixed with sCatch-GFP-ΔCatcher (green) leading to no sfCherry reconstitution (magenta). Scale bars: 20 µm.

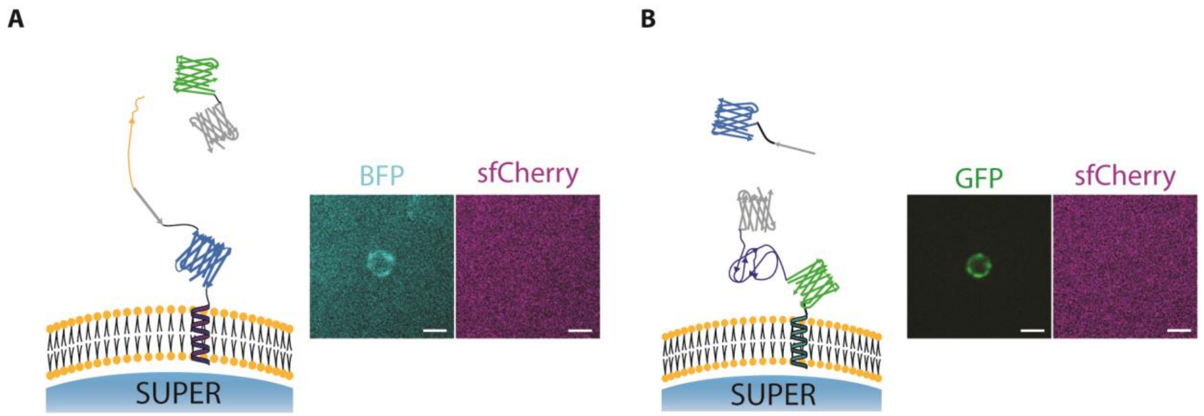


**Figure S2**: SpyTag/SpyCatcher interaction is indispensable for sfCherry reconstitution on SUPER templates. **A)** Representative confocal images of reconstitution of cell-free expressed InterTag-BFP (cyan) on the membrane of 5 µm SUPER templates mixed with sCatch-GFP-ΔCatcher showing lack of sfCherry (magenta) reconstitution. Scale bar: 5 µm **B)** Representative confocal images of reconstitution of cell-free expressed InterCatch-GFP (green) on the membrane of 5 µm SUPER templates mixed with sTag-BFP-ΔTag showing absence of sfCherry (magenta) signal. Scale bar: 5 µm.

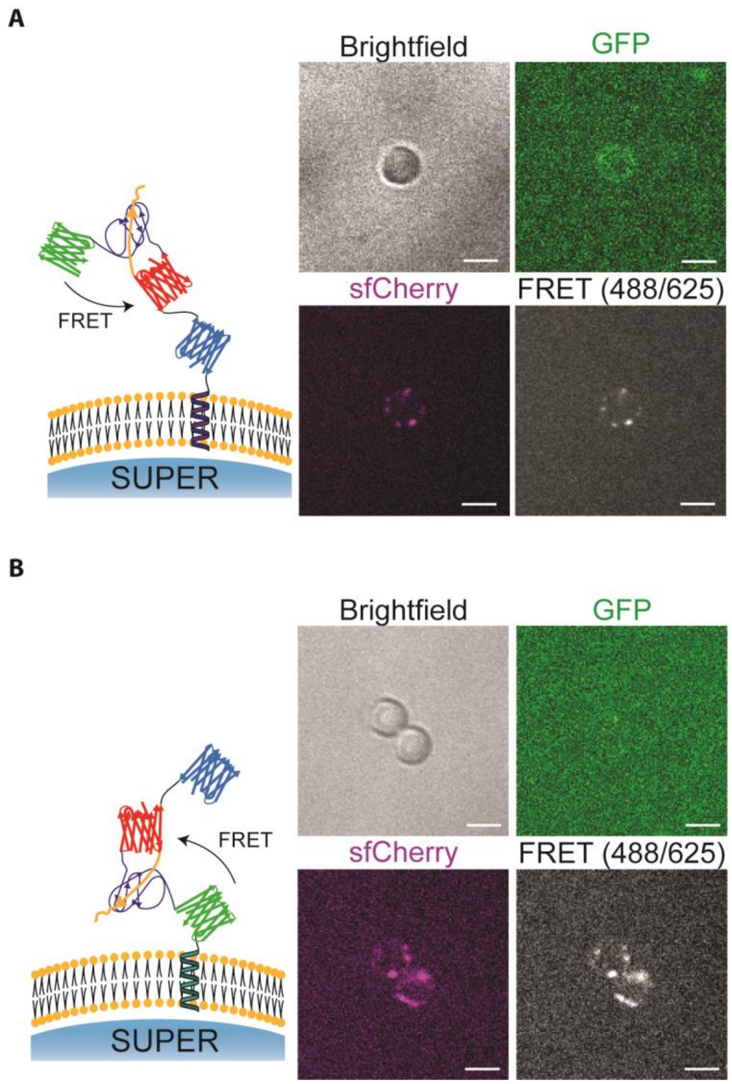


**Figure S3**: Fluorescence resonance energy transfer (FRET) between GFP and sfCherry on the membrane of SUPER templates. **A)** Representative confocal images of reconstitution of cell-free expressed InterTag-BFP on the membrane of 5 µm SUPER templates mixed with sCatch-GFP (green) and forming sfCherry (magenta), leading to FRET between GFP and sfCherry. Scale bar: 5 µm **B)** Representative confocal images of reconstitution of cell-free expressed InterCatch-GFP (green) on the membrane of 5 µm SUPER templates mixed with sTag-BFP and forming sfCherry (magenta), leading to FRET between GFP and sfCherry. Scale bar: 5 µm.

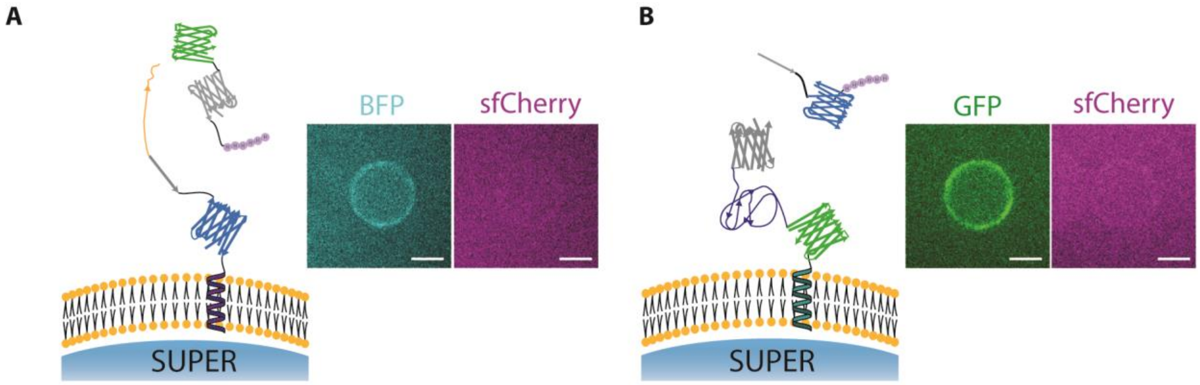


**Figure S4**: SpyTag/SpyCatcher interaction is indispensable for sfCherry reconstitution on 20 µm SUPER templates. **A)** Representative confocal images of bottom-up reconstitution of membrane proteins InterTag-BFP (cyan) on the membrane of 20 µm SUPER templates and no fluorescent sfCherry (magenta) formation when mixed with purified sCatch-GFP-ΔCatcher. Scale bars: 10 µm. **B)** Representative confocal images of bottom-up reconstitution of membrane proteins InterCatch-GFP (green) on the membrane of 20 µm SUPER templates and no functional sfCherry (magenta) formation when mixed with purified sTag-BFP-ΔTag. Scale bars: 10 µm.

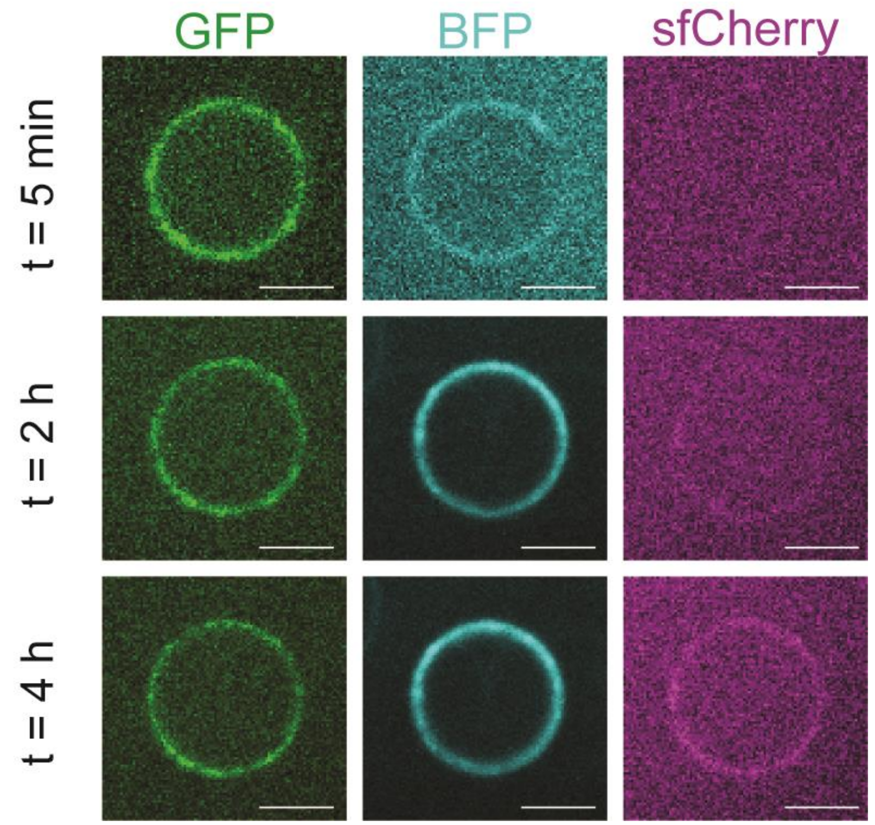


**Figure S5:** Reconstitution of fluorescent sfCherry mediated by SpyTag-SpyCatcher interaction over time. Shown are representative confocal images of cell-free reconstituted membrane protein InterCatch-GFP (green) on SUPER templates and rapid localization of SUVs harboring sTag-BFP (cyan) before sfCherry formation (magenta). Scale bar: 10 µm.

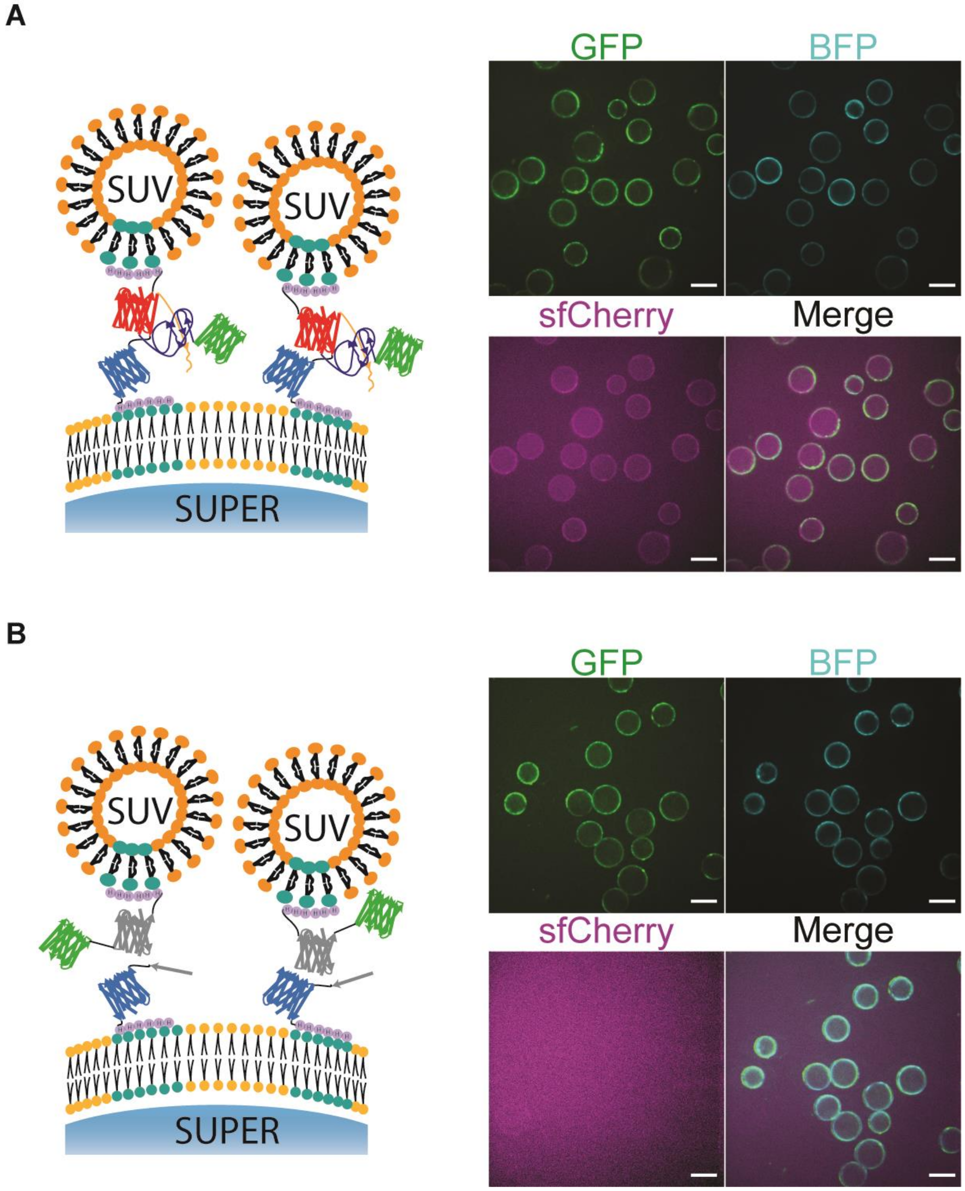


**Figure S6**: sfCherry reconstitution is relied on SpyTag/SpyCatcher bond formation when purified protein fragments create the membrane interface. **A)** Representative cohort images of reconstituted sfCherry (magenta) that fluoresces in the space between SUVs harboring sCatch-GFP (green) and SUPER templates displaying sTag-BFP (cyan). Scale bars: 20 µm. **B)** Representative cohort images of no reconstituted sfCherry (magenta) in the space between SUVs harboring sCatch-GFP-ΔCatcher (green) and SUPER templates displaying sTag-BFP (cyan). Scale bars: 20 µm.

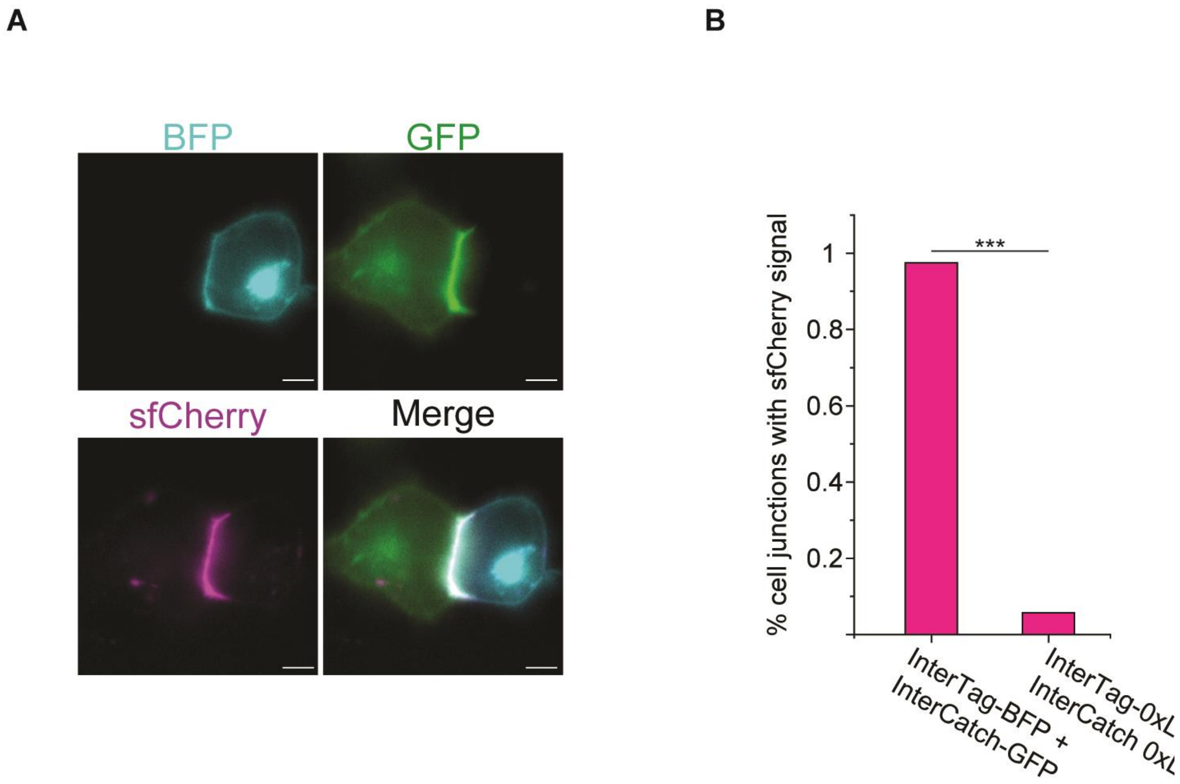


**Figure S7**: **A)** Representative single cell image of sfCherry (magenta) reconstituted through InterTag-BFP (cyan) and InterCatch-GFP (green). Scale bar: 5 µm. **B)** Plot showing percent of cell-cell junctions with reconstituted sfCherry signal for co-cultures involving InterTag-BFP and InterCatch-GFP and between InterTag-0xL and InterCatch-0xL. *** represent $p < 0.001$.

**Table S1.** Total number of cell junctions showing positive or negative fluorescence sfCherry signal in different linker combination groups. IC#:IT# represents co-culture of InterCatch #xL cells with InterTag #xL cells where # denotes the number of GGGGS linkers in each cell line.

| Total number of GGGGS linkers | 0 | 1 | | 2 | | | |
|---|---|---|---|---|---|---|---|
| Combinations | IC0:IT0 | IC1:IT0 | IC0:IT1 | IC2:IT0 | IC1:IT1 | IC0:IT2 | |
| Positive | 14 | 7 | 14 | 1 | 3 | 7 | |
| Negative | 240 | 72 | 90 | 51 | 25 | 31 | |
| Total number of GGGGS linkers | 3 | | | | 4 | | |
| Combinations | IC3:IT0 | IC2:IT1 | IC1:IT2 | IC0:IT3 | IC3:IT1 | IC2:IT2 | IC1:IT3 |
| Positive | 2 | 23 | 61 | 24 | 17 | 100 | 73 |
| Negative | 59 | 30 | 56 | 20 | 23 | 57 | 77 |
| Total number of GGGGS linkers | 5 | | 6 | | | | |
| Combinations | IC3:IT2 | IC2:IT3 | IC3:IT3 | | | | |
| Positive | 164 | 132 | 272 | | | | |
| Negative | 72 | 65 | 100 | | | | |

**Table S2**. Constructs obtained from Addgene.

| | |
|---|---|
| **TfR-sfGFP-myc tag-SpyCatcher003** | https://www.addgene.org/133451/ |
| **pcDNA3.1(+)_SpyCatcher-6aa-sfCherry(1-10)** | https://www.addgene.org/117484/ |
| **pSFFV-SpyTag-sfCherry2(11)-TagBFP** | https://www.addgene.org/117485/ |
| **SpyTag003-mKate2** | https://www.addgene.org/133452/ |
| **pDisplay-LAP2-CFP-TM** | https://www.addgene.org/34842/ |
| **pSBbi-GP** | https://www.addgene.org/60511/ |
| **pSBbi-BP** | https://www.addgene.org/60512/ |
| **pCMV(CAT)T7-SB100** | https://www.addgene.org/34879/ |

**Table S3**. Primers used and constructs generated in this study

<u>Oligos</u>

- 1x GGGGS: 5’ GCTCTTCccgcgggaggcggtggatcggaacaGAAGAGC 3’
- 2x GGGGS: 5’ GCTCTTCccgcgggaggcggtggatcgggaggcggtggatcggaacaGAAGAGC 3’
- 3x GGGGS: 5’GCTCTTCccgcgggaggcggtggatcgggaggcggtggatcgggaggcggtggatcggaacaGAAGAGC 3’

| Construct (Cloning method) | Component | Primer | Template plasmid |
|---|---|---|---|
| **0xSBCatch (Gibson)** | TfR | TfR_fwd: ccaagctggcctctg*gccaccatg*GATCAAGCTAGATCAGCATTCTC<br><br>TfR_rev: gtttttgttcATAGCCCAAGTAGCCAATCATAAATC | **TfR-sfGFP-myc tag-SpyCatcher003** |
| | SpyCatcher003 + sfC1-10 | C3_sfC110_fwd: tgggctatGAACAAAAACTCATCTCAGAAGAG<br><br>C3_sfC110_rev: caagcttggcctgacCTAGTCCTCGTTGTGGCTGG | **TfR_sfGFP_sC3_sfC110_Zeo** |
| | SB backbone (GFP) | SfiI digestion | **pSBbi-GP** |
| **0xSBTag**<br><br>**(Gibson)** | IgK + SpyTag003 + sfC11 | Igk_T3_sfC11_xTag_fwd: accccaagctggcctctg*gccaccatg*GAGACAGACACACTC<br><br>CTGCTATGG<br><br>Igk_T3_sfC11_xTag_rev: ttgttcggcGGTGCTGTGTCTGGCCTCGG | **pdgfr_BFP_sfc11_sT3_Zeo** |
| | PDGFR + Myc | pdgfr_xTag_fwd: cacagcaccGCCGAACAAAAACTCATCTC | **pdgfr_BFP_sfc11_sT3_Zeo** |

| | | | |
|---|---|---|---|
| | | pdgfr_xTag_rev: ccccaagcttggcctgacCTAACGTGGCTTCTTCTGCC | |
| | SB backbone (BFP) | SfiI digestion | **pSBbi-BP** |
| **1x,2x,3x SBCatch**<br><br>**(Golden Gate)** | GGGGS Oligo | SapI digestion + **0xSBCatch** | |
| **1x,2x,3xSBTag**<br><br>**(Golden Gate)** | GGGGS Oligo | SapI digestion + **0xSBTag** | |
| **3xSB ΔTag (Golden Gate)** | | 3xSB_NoTag_SapI_fwd: ttttttGCTCTTCaggctacaccatcgtgga<br><br>3xSB_NoTag_SapI_rev: ttttttGCTCTTCagccgtcaccagtggaacctg | **3xSBTag** |
| **pT7-CFE1-InterCatch-GFP (Gibson)** | Vector | BB-FWD: CACCACCACCACTAATAAAGATC<br><br>BB-REV: CATATTATCATCGTGTTTTTCAAAGG | **pT7-CFE1-6xHis-HA** |
| | TfR-sfGFP-SpyCatcher003-sfCherry1-10 | xCatch-FWD: TTTTCCTTTGAAAAACACGATGATAATATGATGATGGATCAAGCTAGATCAGC<br><br>xCatch-REV: TCAGTCAGATCTTTATTAGTGGTGGTGGTGCTAGTCCTCGTTGTGGCTGG | **TfR-sfGFP-SpyCatcher003-sfCherry(1-10)** |
| **pT7-CFE1-InterTag-BFP (Gibson)** | Vector | BB-FWD: CACCACCACCACTAATAAAGATC | **pT7-CFE1-6xHis-HA** |

| | | | |
|---|---|---|---|
| | | BB-REV: CATATTATCATCGTGTTTTTCAAAGG | |
| | SpyTag003-sfCherry11-tagBFP-PDGFR | xTag-FWD: TTTTCCTTTGAAAAACACGATGATAATATGCGTGGCGTTCCTCATATTG<br><br>xTag-REV: TCAGTCAGATCTTTATTAGTGGTGGTGGTGCTAACGTGGCTTCTTCTGC | **IgK-SpyTag003-sfCherry11-tagBFP-PDGFR** |
| **pET28b-sCatch-GFP (Gibson)** | Vector | BB-FWD: CACCACCACCACCACCAC<br><br>BB-REV: AAAAAACCTCCTTACTTTCTAGTCTCAAG | **pET28b_(R5Q5)(R5Q6)F20_His_Lys_RBS** |
| | sfGFP-SpyCatcher003-sfCherry1-10-6xHis | sCatch-FWD: TCTTGAGACTAGAAAGTAAGGAGGTTTTTTATGCGTAAAGGCGAAGAGC<br><br>sCatch-REV: AGCCGGATCTCAGTGGTGGTGGTGGTGGTGGTCCTCGTTGTGGCTGGTGATG | **pT7-CFE1-InterCatch-GFP** |
| **pET28b-sTag-BFP (Gibson)** | Vector | BB-FWD: CACCACCACCACCACCAC<br><br>BB-REV: AAAAAACCTCCTTACTTTCTAGTCTCAAG | **pET28b_(R5Q5)(R5Q6)F20_His_Lys_RBS** |
| | SpyTag-sfCherry11-tagBFP-6xHis | sTag-FWD: TCTTGAGACTAGAAAGTAAGGAGGTTTTTTatgcgtggcgttcctcatattg<br><br>sTag-REV: AGCCGGATCTCAGTGGTGGTGGTGGTGGTGCAGATCCTCTTCTGAGATGAG | **pT7-CFE1-InterTag-BFP** |

| | | | |
|---|---|---|---|
| **pET28b-sCatch-GFP-ΔCatcher (Gibson)** | Vector | BB-FWD: CACCACCACCACCACCAC<br><br>BB-REV: AAAAAACCTCCTTACTTTCTAGTCTCAAG | **pET28b_(R5Q5)(R5Q6)F20_His_Lys_RBS** |
| | sfGFP-sfCherry1-10-6xHis | sCatch-dC-FWD: TCTTGAGACTAGAAAGTAAGGAGGTTTTTTatgcgtaaaggcgaagagc<br><br>sCatchdC-REV: AGCCGGATCTCAGTGGTGGTGGTGGTGGTGGTCCTCGTTGTGGCTGGTGATG | **pT7-CFE1-InterCatch-GFP** |
| **pET28b-sTag-BFP-ΔSpyTag (Gibson)** | Vector | BB-FWD: CACCACCACCACCACCAC<br><br>BB-REV: AAAAAACCTCCTTACTTTCTAGTCTCAAG | **pET28b_(R5Q5)(R5Q6)F20_His_Lys_RBS** |
| | sfCherry11-tagBFP-6xHis | sTagdT-FWD: TCTTGAGACTAGAAAGTAAGGAGGTTTTTTATGTACACCATCGTGGAGCAGTAC<br><br>sTagdT-REV: AGCCGGATCTCAGTGGTGGTGGTGGTGGTGCAGATCCTCTTCTGAGATGAG | **pT7-CFE1-InterTag-BFP** |
| **pT7-CFE1-sCatch-GFP (Gibson)** | Vector | BB-FWD: CACCACCACCACTAATAAAGATC<br><br>BB-REV: CATATTATCATCGTGTTTTTCAAAGG | **pT7-CFE1-6xHis-HA** |
| | sfGFP-SpyCatcher003-sfCherry1-10 | sCatch-FWD: TTTTCCTTTGAAAAACACGATGATAATATGATGCGTAAAGGCGAAGAGC<br><br>sCatch-REV: TCAGTCAGATCTTTATTAGTGGTGGTGGTGCTAGTCCTCGTTGTGGCTG | **pT7-CFE1-InterCatch-GFP** |

| | | | |
|---|---|---|---|
| **pT7-CFE1-sTag-BFP (Gibson)** | Vector | BB-FWD: TAGCACCACCACCACTAATAAAG<br><br>BB-REV: CATATTATCATCGTGTTTTTCAAAGG | **pT7-CFE1-6xHis-HA** |
| | SpyTag003-sfCherry11-tagBFP | sTag-FWD: TTTTCCTTTGAAAAACACGATGATAATATGCGTGGCGTTCCTCATATTG<br><br>sTag-REV: GTCAGATCTTTATTAGTGGTGGTGGTGctaCAGATCCTCTTCTGAGATGAG | **pT7-CFE1-InterTag-BFP** |
| **pT7-CFE1-sCatch-GFP-ΔCatcher (Gibson)** | Vector | BB-FWD: CACCACCACCACTAATAAAGATC<br><br>BB-REV: GGATCCACCCGAGCCA | **pT7-CFE1-6xHis-HA** |
| | sfGFP-sfCherry1-10 | sCatch-dC-FWD: gatctgggttccggtggctcgggtggatccggatctagcggatctatggagg<br><br>sCatch-dC-REV: TCAGTCAGATCTTTATTAGTGGTGGTGGTGCTAGTCCTCGTTGTGGCTG | **pT7-CFE1-InterCatch-GFP** |
| **pT7-CFE1-sTag-BFP-ΔTag (Gibson)** | Vector | BB-FWD: TAGCACCACCACCACTAATAAAG<br><br>BB-REV: CATATTATCATCGTGTTTTTCAAAGG | **pT7-CFE1-6xHis-HA** |
| | sfCherry11-tagBFP | sTag-dT-FWD: TTTTCCTTTGAAAAACACGATGATAATATGTACACCATCGTGGAGCAG<br><br>sTag-dT-REV: GTCAGATCTTTATTAGTGGTGGTGGTGctaCAGATCCTCTTCTGAGATGAG | **pT7-CFE1-InterTag-BFP** |

**Table S4.** Calculated p-values for comparing statistical significance between groups in **Figure 5D**

| | 0x | 1x | 2x | 3x | 4x | 5x | 6x |
|---|---|---|---|---|---|---|---|
| 0x | | 0.654 | 1 | 3.35E-21 | 1.11E-39 | 1.86-64 | 7.59E-70 |
| 1x | | | 1 | 1.57E-10 | 1.55E-22 | 7.22E-40 | 1.48E-44 |
| 2x | | | | 4.83E-9 | 2.78E-18 | 3.08E-31 | 3.39E-35 |
| 3x | | | | | 0.008 | 3.11E-12 | 4.61E-16 |
| 4x | | | | | | 0.0017 | 5.32E-6 |
| 5x | | | | | | | 1 |
| 6x | | | | | | | |
